# An algorithmic framework for isoform-specific functional analysis

**DOI:** 10.1101/2022.05.13.491897

**Authors:** Guy Karlebach, Leigh Carmody, Jagadish Chandrabose Sundaramurthi, Elena Casiraghi, Peter Hansen, Justin Reese, Chris J Mungall, Giorgio Valentini, Peter N. Robinson

## Abstract

Gene Ontology (GO) overrepresentation analysis characterizes the biological mechanisms common to sets of differentially expressed genes identified by high-throughput experiments. To date, GO overrepresentation analysis has mainly been used to evaluate differentially expressed genes, but short- and long-read RNA-seq technologies now allow increasingly accurate identification of differential alternative splicing. The function of most splice isoforms remain unknown, but if acccurate predictions could be made, overrepresentation analysis could be applied to differentially spliced isoforms to assess the functional implications of alternative splicing in RNA-seq experiments. We present *isopret* (Isoform Interpretation), a new paradigm for isoform function prediction based on the expectation-maximization framework. *isopret* leverages the relationships between sequence and functional isoform similarity to infer isoform specific functions in a highly accurate fashion. This enabled us to adapt GO overrepresentation analysis, which to date has been limited to differential gene expression, to be extended to assess overrepresentation of GO annotations in differentially spliced isoforms. An analysis of 100 RNA-seq studies including investigations of development, cancer, and common disease demonstrated that expression and splicing regulate different sets of biological functions. We make *isopret* predictions freely available in a desktop application that can be used to analyze differential expression and splicing in any bulk RNA-seq dataset.

## Introduction

Gene Ontology (GO) overrepresentation analysis of differentially expressed gene sets has been widely used to gain insight into the biological mechanisms. The two major components of the GO project are the ontology itself as well as the GO annotations. For instance, the *NF1* gene product neurofibromin is annotated to GO terms including *positive regulation of GTPase activity* (GO:0043547). The assumption of GO overrepresentation analysis is that if the set of genes identified by an experiment (e.g., the set of differentially expressed genes) contain more genes annotated to a given GO term than expected by chance, then the term is likely to be linked to an important biological driver or attribute underlying the experimental results.[1]

To date, GO overrepresentation analysis has predominantly been used to evaluate differentially expressed genes.[2, 3] However, the advent of accurate and low-cost short- and long-read RNA sequencing (RNA-seq) technology now offers the opportunity to study not only differential gene expression but also differential alternative splicing.[4] More than 90% of human genes undergo alternative splicing, a process that involves alternative patterns of intron removal to generate alternatively spliced transcripts that differ in their coding capacity, stability, or translational efficiency.[5] Until recently, a typical assumption of biomedical analysis was that the functions of canonical isoforms provide a faithful representation of the function of a gene or a gene product.[6] However, alternatively spliced isoforms provide functional diversity at the level of enzymatic activities and subcellular localizations, as well as protein-protein, protein-DNA, and protein-ligand physical interactions.[7] Although individual examples of functional differences between isoforms have been confirmed experimentally, the function of most splice isoforms remain unknown.[8] Additionally, but it remains largely unknown how expression and splicing are integrated to implement global cellular processes. In principle, there are two possibilities: (1) differential alternative splicing (DAS) and differential gene expression (DGE) could be regulated simultaneously to mediate similar biological functions; or (2) DAS and DGE could be regulated separately to achieve distinct sets of biological goals.

RNA-seq technologies now allow isoform expression to be measured with steadily increasing accuracy and comprehensiveness.[4, 9] However, the computational resources needed to address questions of global functional impact of DAS and DGE have been lacking. The problem is thus twofold: Methods to assess DGE and DAS in RNA-seq data are essential to identify the impact of these processes in data from high-throughput experiments. Secondly, changes in biological function owing to both DGE and DAS need to be understood. In particular, it is highly desirable to obtain comprehensive GO annotations at the isoform level to enable isoform-focused GO overrepresentation analysis. In this work, we model the relationships between genes, isoforms and functions (GO annotations) as a graph and formulate the isoform function assignment problem as a global optimization problem. Based on this, we use expectation maximization to derive GO annotations for different isoforms. In addition, we provide software that can be used to visualize results of GO overrepresentation analyzes for both differentially expressed genes and differentially spliced isoforms. On datasets from a broad set of conditions including cancer, autoimmunity, and neurodegenerative disease, we demonstrate that DGE and DAS tend to have distinct and characteristic patterns of overrepresented GO terms.

## Results

Gene Ontology (GO) resources traditionally have described the function of gene products without distinguishing the potentially different functions of specific isoforms. Here, we present a method for inferring GO annotations for isoforms based on patterns of gene-level annotations across the genome and leverage these annotations to implement an algorithm to assess isoform-specific GO overrepresentation analysis. For the sake of brevity, we will use the expressions “GO term” and “function” interchangeably.

### Assignment of GO annotations to isoforms using expectation-maximization

The functions of proteins are largely determined by their amino-acid sequence. An accurate and complete mapping from sequence to function is not currently possible, however, the degree to which functions are shared by a pair of proteins can be estimated from their sequence similarity. Based on the assumption that the functions of an isoform represent a subset of the functions of the associated gene, we developed an optimization procedure that maximizes the agreement between functional similarity and sequence similarity.

More specifically, our approach makes the following assumptions: 1. The GO terms assigned to a gene correspond to the function of at least one isoform of the gene. 2. Isoform pairwise sequence similarity tends to be positively correlated with the number of GO annotations they share. 3. The pairwise isoform sequence similarity scores for different isoform pairs can be regarded as independent random variables if the genes to which encode the isoforms in each pair share at least one GO term.

Our approach is based on expectation-maximization (EM), which means that we alternate between two optimization steps until convergence is reached. First a matrix of pairwise local alignment scores is calculated for all pairs of isoforms. This matrix only needs to be calculated once. Then, a model *M* is defined that distributes the GO annotations of a gene to the gene’s isoforms. A quadratic regression equation is defined that predicts the local alignment score of a pair of isoforms on the basis of the number of shared GO terms.

As a starting point for the optimization, our method distributes the gene-level annotations to coding isoforms according to their predicted InterPro domains and the GO terms associated with these domains. GO annotations not related to InterPro domains are initially not assigned to any isoform (Fig. 1a). InterPro [10] is a database that stores amino acid signatures that can be matched to protein sequences using regular expressions or Hidden Markov Models. An isoform-by-isoform matrix is constructed whereby cell (*i, j*) contains the normalized local sequence alignment score for isoforms *i* and *j* (Fig. 1b). These values constitute the dependent variable that will be predicted from the number of shared GO terms. For the independent variable values, an isoform-by-GO term matrix is constructed with a 1 in cell (*i, j*) if isoform *i* is annotated to GO term *j* and otherwise 0 (Fig. 1c). Multiplication of this matrix by its transpose yields an isoform-by-isoform matrix where each cell contains the number of shared GO annotations. Our model posits a quadratic relationship between the number of shared GO annotations and the sequence similarity (Figure S1A) and attempts to minimize the error term (sum of squared residuals) by iteratively adjusting the assignment of GO terms to isoforms using a genetic algorithm, and then adjusting the parameter values for the quadratic model. This is achieved using an EM framework (Methods) consisting of the following two steps:

**Figure 1.**
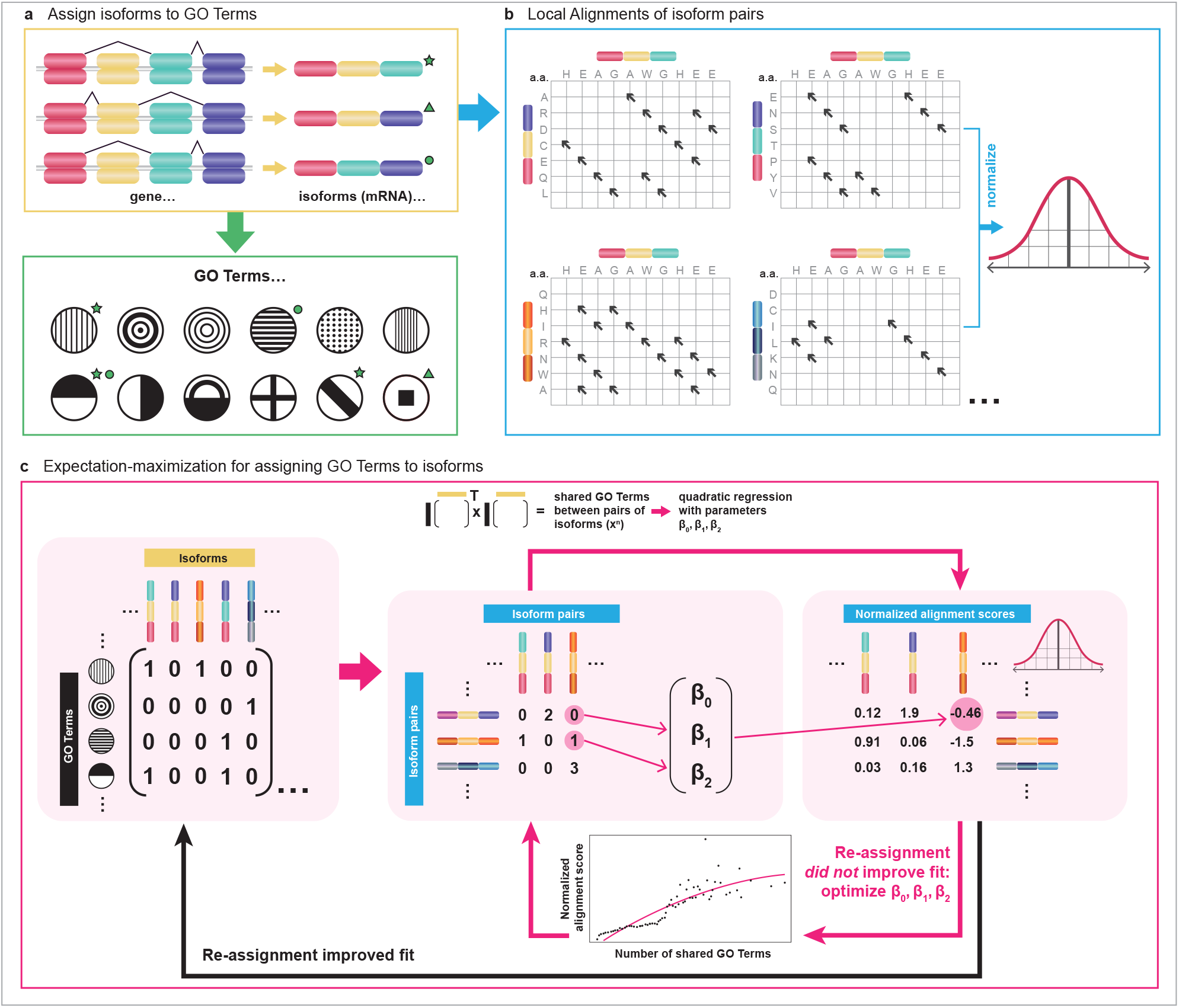
Optimization algorithm for assigning functions from genes to isoforms. A) Assignments to isoforms are initialized with subset of their gene’s GO terms on the basis of their predicted InterPro domains. B) Local alignment scores are computed between every pair of isoforms, and the scores are log transformed and standardized. C) The isoform-to-GO-term binary matrix is multiplied by its transpose to obtain a matrix of the number of shared GO terms between each pair of isoforms. These values are then used as the independent variable in a quadratic model with parameters *β*_0_, *β*_1_, *β*_2_ to predict the normalized local alignment scores between pairs of isoforms. The GO term assignment is optimized by a stepwise procedure that uses a genetic algorithm (Methods) until no further improvement in the model fit is possible (black arrow), at which point the GO term assignment is fixed and new parameter values *β*_0_, *β*_1_, *β*_2_ are obtained by optimizing the fit of the model (pink arrow). These two steps of GO term assignment optimization and parameter values optimization are repeated consecutively until no further improvement can be obtained.

**E step** Find the best assignments of the GO terms for a model *M* that predicts the pairwise alignment scores between isoforms on the basis of the number of their shared GO terms

**M step** Optimize the parameters of the regression equation using the best GO terms assignments of the E step.

The *isopret* algorithm was used to generate function predictions for 85,617 isoforms of 17,900 protein-coding genes. For 17,401 of these genes (97.2%), *isopret* inferred isoform specific functions.

### Consistency with InterPro Domain Predictions

One measure by which one can assess the accuracy of isoform function prediction is by its consistency with the domains provided by InterPro. Some of the InterPro domains have been associated with GO terms via interpro2GO, which is a semi-automatic annotation system that maps InterPro entries to GO terms in one-to-one manner.[10] Interpro2GO predicts that a protein has a given GO annotation if the corresponding protein is predicted to have a certain domain, which can correspond for example to an active site or a motif characteristic of a certain protein family. Therefore, a function assignment that is consistent with protein primary structure implies that the more GO terms shared by two protein sequences, the more InterPro domains they will also share. To evaluate how reliably this relationship is expressed in our isoform function assignments, we determined the number of shared InterPro domains as a function of the number of shared GO terms, when the latter are assigned using *isopret* and by assigning an isoform with all of its gene’s terms. Fig. 2a displays the changes in the number of shared InterPro domains as the number of shared GO terms increases for these two assignment methods. As can be seen in the figure, there is a sharper increase for the *isopret* assignments, in contrast to the gene-level assignments which do not result in a steady increase in the median number of shared domains. The improvement in correlation is reflected by the values of Kendall’s tau rank correlation coefficient, which were 0.38 for the Isopret assignment and 0.18 for gene-level assignment, whereby higher values of *τ* indicate better agreement of the order of ranks between isoform functions and InterPro domains, with 0 at no agreement and 1 at perfect agreement of ranks. The figure was generated using a sample of unique isoform pairs containing 50,000 Interpro domains.

**Figure 2.**
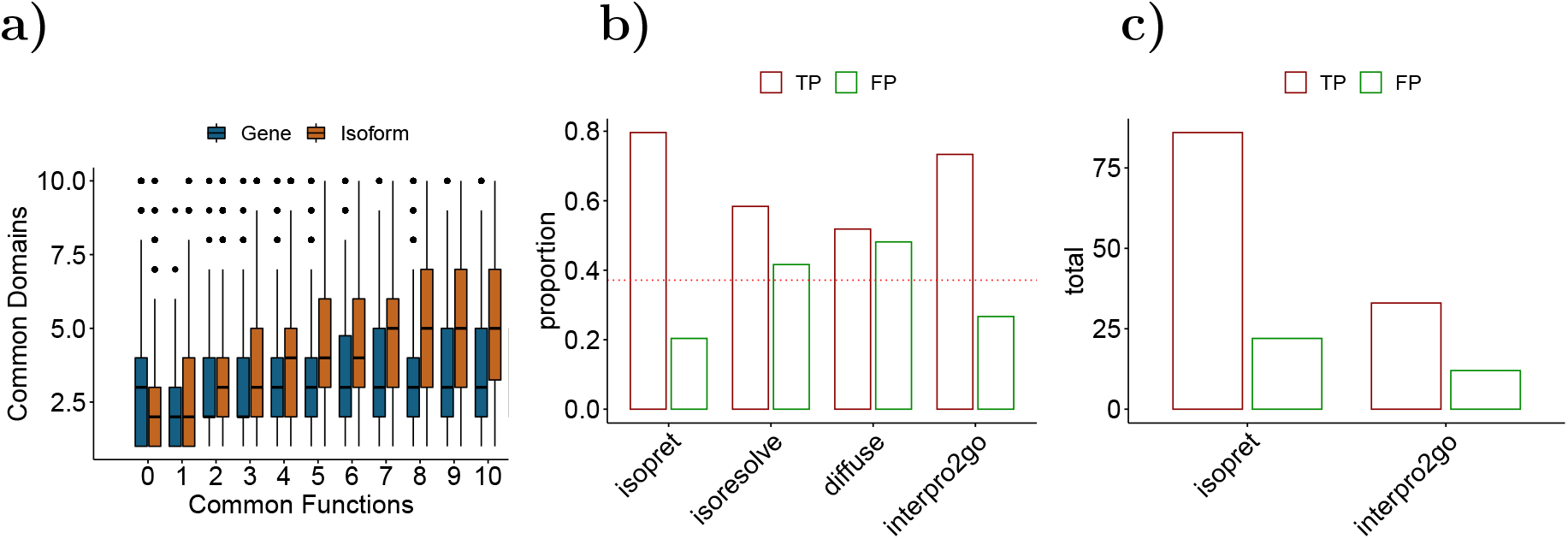
Isoform-specific GO annotation predictions. **a)** Number of shared InterPro domains as a function of the number of shared GO terms. Brown: Boxplots for the number of shared InterPro domains between isoforms that share at least one domain (y-axis) as a function of different numbers of shared GO terms that are assigned by the *isopret* algorithm (x-axis), from 0 to 10 shared GO terms. The median of the number of shared InterPro domains increases with the number of shared GO terms. Blue: The same boxplots for the number of shared InterPro domains, for different numbers of shared GO terms, where all the GO terms of a gene are assigned to all of its isoforms. While the Isopret-assigned GO terms are selected from the gene’s GO terms and Interpro2GO, the correlation between the count of shared GO terms and the count of shared InterPro domains is higher for the former (Kendall’s tau 0.38 vs 0.18, respectively). **b)** Comparison of the proportion of correct GO term assignments and incorrect assignments using a gold-standard of curated 307 isoform functions. The horizontal red dotted line marks the proportion of negative examples in the test set. **c)** Comparison of the number of correct GO term assignments and incorrect assignments using a gold-standard of curated isoform functions.

### Comparison to existing annotations and prediction methods

Many existing methods for isoform function prediction leverage multiple-instance learning (MIL) in different ways. MIL is a supervised learning approach in which the learner receives a set of labeled bags that contain multiple instances. In the case of isoform prediction, each gene is considered to be a bag of alternatively spliced isoforms. In order to predict GO annotations, the MIL model assumes that at least one of the isoforms must be annotated (i.e., a positive example) and attempts to optimize a model based on similarity of attributes including isoform-specific co-expression, isoform sequence data, proteomics data, and distribution of protein motifs in isoforms.[11, 12, 13, 14, 15, 16] Newer studies have adapted Bayesian logistic regression, domain adaptation and deep learning. We compared our method to previously published methods whose authors provided code or downloadable files (Supplemental Table S1), namely, IsoResolve, an approach based on domain adaptation which was shown to outperform other available methods by its authors,[17] DIFFUSE,[16] which is a deep-learning approach, and interpro2GO.

In order to be able to compare different GO term assignment methods, we curated 307 isoform specific functions from 72 publications, including a total of 149 different isoforms from 62 genes annotated to 97 different GO terms (Supplementary Material). Since approximately 40% of our curated examples are negative labels, we expect a false positive rate of less than 40% and a true positive rate of more than 60% from any method that surpasses random assignment (Fig. 2b). As can be seen, only *isopret* and interpro2GO perform better than random assignment. In addition, *isopret* has a higher rate of true positives and a lower rate of false positives, with TP/FP rates of 0.8/0.2 and 0.73/0.27 for isopret and interpro2GO, respectively. To further compare *isopret* assignments to those derived from interpro2GO, we examined the counts of true and false positives (Fig. 2c). As the figure illustrates, *isopret* predicts more than twice as many terms as interpro2GO (108 vs. 45) with better accuracy. Therefore, for the remaining analyses in this paper we will use *isopret* GO term assignments.

### Isoform-specific GO overrepresentation analysis

Gene-level GO overrepresentation analysis compares the proportion of genes annotated to a given GO term in a set of differentially expressed (or otherwise selected) genes as compared to the proportion of genes in the population (either all genes, all genes with at least one sequence read in an RNA-seq experiment, or other similar definitions) that are annotated to the term. Prior to analysis, annotations are propagated to parent terms according to the true-path rule.[1] Analysis of overrepresentation can be performed by a Fisher exact test (FET) or related procedures such as parent-child analysis.[18]

To perform isoform GO analysis and compare its results to gene-level GO overrepresentation analysis, one must first identify differentially expressed genes and differentially spliced isoforms in a dataset. We have previously developed HBA-DEALS, which applies hierarchical Bayesian analysis to characterize differential expression and splicing in the same analysis.[19] We used this tool in the analysis presented here but any approach that provides such sets of genes and isoforms could be substituted.

For detection of differential alternative splicing enriched terms, our analysis takes as input the list of isoforms characterized as differential and compares their GO annotations with those of the population of all isoforms with at least one called RNA-seq read in the dataset. If certain GO terms occur more often than would be expected by chance, they are flagged as enriched terms (Methods).

### Characterization of DGE and DAS in a collection of 100 RNA-seq experiments

In order to investigate profiles of overrepresented GO terms in sets of differentially expressed genes (DGE) and sets of differentially spliced isoforms (DAS), we assembled a collection of 100 RNA-seq experiments with a variety of focus areas including disease pathophysiology, cancer, infectious disease, as well as cell and molecular biology (Supplemental Tables S2-S5). Sequenced reads were processed with fastp for quality control and mapped by STAR, and isoforms were quantified with RSEM (Methods). DGE and DAS were called using HBA-DEALS.[19] In order to assess to what extent differential expression and differential splicing are associated with the same GO terms (and thus by our hypothesis affect the same or different biological functions), we computed the Jaccard index for the sets of GO terms enriched with the sets of differentially expressed genes and differentially spliced isoforms in each data set. (Fig. 3a; Methods). 75% of all indices are less than 0.06, indicating that DGE and DAS are generally associated with different GO terms. Most enriched GO terms appeared as either DAS or DGE (Fig. S6). In general, there were more overrepresented GO terms for DGE than for DAS (Fig. 3b).

**Figure 3.**
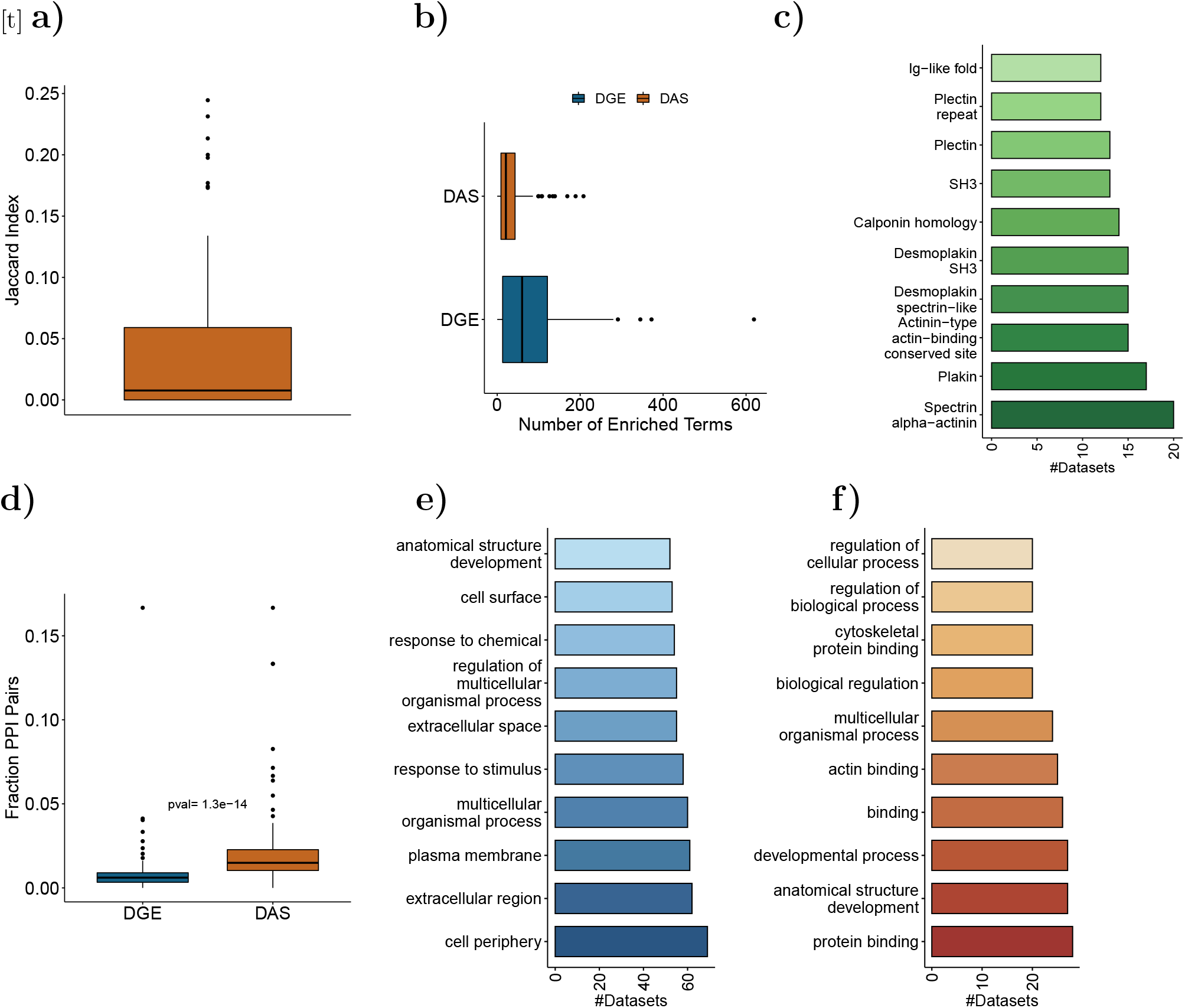
Comparison of DAS and DGE genes in 100 RNA-seq experiments. **a)** Overlap of DGE and DAS enriched GO terms. The Jaccard index between GO terms enriched in DGE and DAS, computed over 100 RNA-Seq datasets. **b)** Box plots of the number of GO terms in DGE- and DAS-enriched per dataset. The most common enriched InterPro domains. **d)** Proportion of PPI interacting genes. The proportions of gene pairs that have a PPI in the BioGrid database, for pairs of genes that are DGE and pairs of genes that have DAS isoforms. **e)** Most common DGE-enriched GO terms. **f)** Most common DAS-enriched GO terms.

We additionally performed overrepresentation analysis for InterPro domains affected by DAS, comparing domains in differentially spliced isoforms with the population of all isoforms (Fig. 3c; Methods). The mean number of enriched domains per dataset was 21.5, and the median was 18. The 10 most common enriched domains are displayed in Figure 3c. Co-enrichment of protein domains in the same dataset is correlated with the frequency of their co-occurrence on the same isoforms, where the former is higher than the latter (Figure S4). This is possibly due to the fact that co-occurrence of a domain in the same isoform implies a functional relationship that can be utilized when the domains appear on different isoforms.

We calculated the proportion of gene pairs that undergo DGE and share a PPI and the proportion of genes that have at least one DAS isoform and share a PPI (Fig. 3d, Methods). The proportion was more than twofold larger for DAS (*p* = 1.3·10^*−*14^, Mann-Whitney test). This confirms and extends previously reported results on enrichment of DAS isoforms involved in PPI in cancer.[20]

Several of the most common DGE-enriched GO terms were related to extracellular functions or signals, for example *cell surface* (GO:0009986), *extracellular region* (GO:0005576), and *response to chemical* (GO:0042221) (Fig. 3e). In contrast, several of the most common DAS-enriched GO terms were related to protein-protein interactions, for example *actin binding* (GO:0003779), *cytoskeletal protein binding* (GO:0008092), and *protein binding* (GO:0005515) (Fig. 3f).

In the following text, we show how the *isopret* predictions can be used with the isopret-gui tool to make testable hypotheses about the role of differential alternative splicing in biological or medical research contexts. The isopret-gui tool takes the output file of HBA-DEALS[19] as input, but calls from other analysis methods could be easily formatted in the same simple, four column file format. The following examples were chosen from the 100 experiments.

Our analysis of RNA-seq data from a comparison of aortic tissue from individuals with abdominal aortic aneurysm and controls showed that the versican gene (*VCAN*) is more highly expressed in patients (fold-change 3.81). 9 *VCAN* isoforms were identified, and three isoforms, including two coding isoforms, were found to be differentially spliced. The longer coding isoform, ENST00000512590, is predicted by *isopret* to have the function *calcium ion binding* (GO:0005509), while the short isoform, ENST:00000513984, is not. This prediction is plausible, since only the longer isoform codes for two calcium-binding epidermal growth factor domains. Additionally, the longer but not the shorter isoform is predicted to be annotated to *extracellular matrix* (GO:0031012). Versican is known to accumulate in aortic aneurysm and altered versican splicing has been observed in abdominal aortic aneurysm.[21, 22] Versican can have intracellular location in development and some forms of cancer,[23, 24] but a specific role for alternative VCAN splicing in altering the balance between intra- and extracellular versican in aortic aneurysms has not been previously described (Figure 4a).

**Figure 4.**
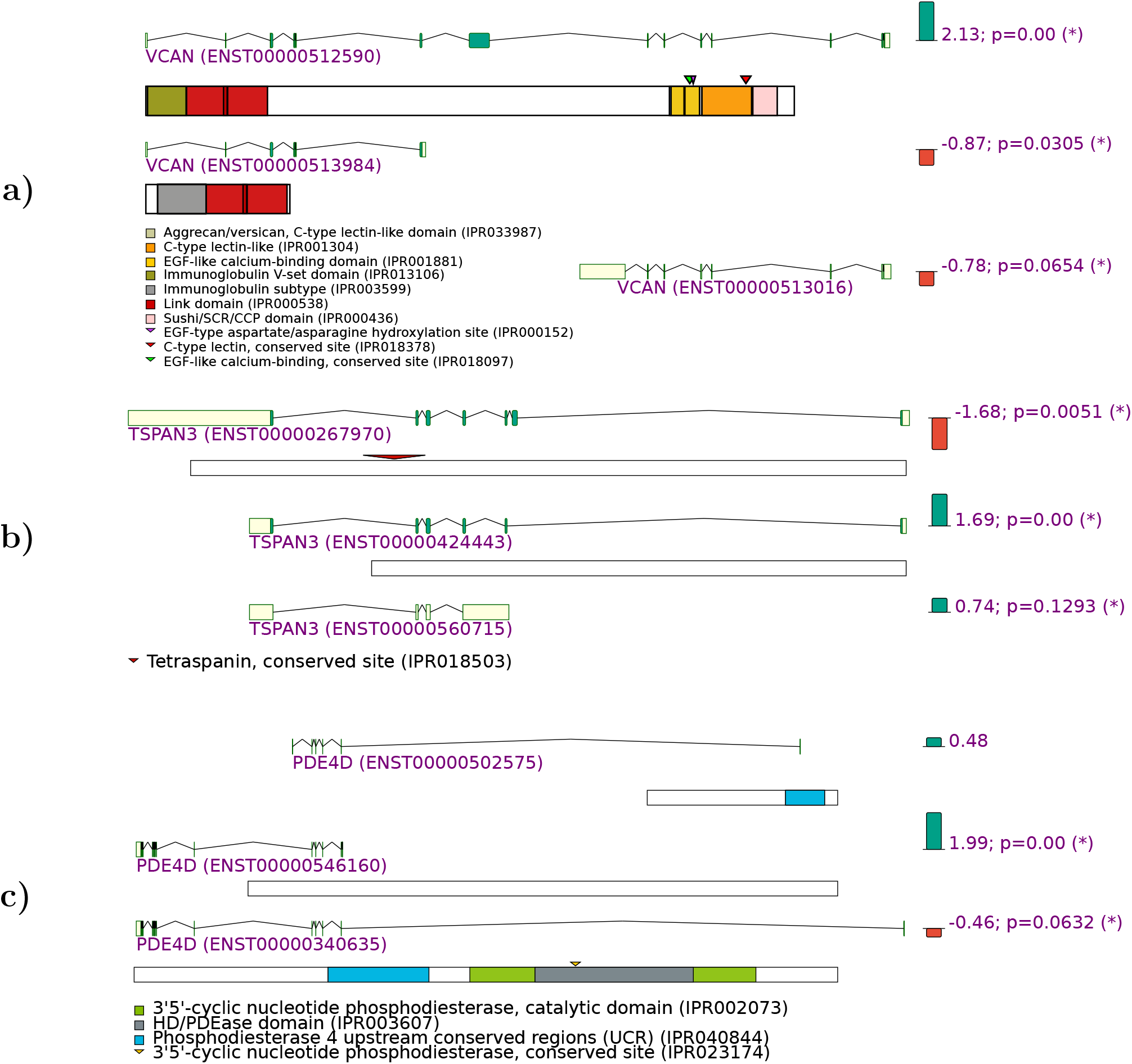
Examples of predicted isoform-specific GO annotations. Transcripts are shown with green boxes representing coding sequences, yellow boxes representing untranslated regions, and inclined lines representing introns. For coding isoforms, a cartoon representing the protein structure and domains is shown. The graphics were exported as SVG files from the isopret-gui app and simplified for display in this Figure. **a)** Versican (*VCAN*) shows DGE in aortic aneurysm (fold-change of overall gene is 3.812 increased in cases compared to controls)). Three isoforms are differentially spliced including a longer and a shorter protein-coding isoform (coding exons are shown in green, untranslated exons in yellow). Only the upregulated isoform is predicted to have calcium binding activity by *isopret*. **b)** Differential *TSPAN3* isoforms. *TSPAN3* gene expression was not differential. One differential isoform with reduced proportionality contains a tetraspanin conserved site. One isoform with increased proportionality does not contain the site, and one is non-coding. **c)** Differential *PDE4D* isoforms. *PDE4D* gene expression was not differential. Only the downregulated isoform codes for the phosphodiesterase catalytic domain.

In a dataset of gastrointestinal stromal tumors compared to adjacent control tissue,[25] the tetraspanin 3 gene (*TSPAN3*) was not differentially expressed, but there was a shift away from an isoform that is predicted by *isopret* for to GO annotation *extracellular exosome* (GO:0070062). This appears plausible because only this isoform contains a tetraspanin conserved site that is located between the second and third transmembrane domains of tetraspanin 3 (Fig. 4b). Tetraspanins play a number of physiological roles in exosomes.[26] Interestingly, exosomes have been shown to confer resistance to chemotherapy in gastric cancer,[27] so our prediction could motivate the hypothesis that alternative splicing alters the transmembrane architecture of tetraspanin 3 and thereby reduces its overall localization in exosome membranes, which could conceivably influence the physiology of exosomes in cancer.

In an experiment in which tacrolimus-treated beta cells were compared with untreated control cells,[28] no differential expression of the phosphodiesterase 4D gene (*PDE4D*) was detected. Only one differentially spliced *PDE4D* isoform (ENST:00000340635) was predicted by *isopret* to have *3’,5’-cyclic-AMP phosphodiesterase activity* (GO:0004115). This prediction is plausible because only one of the three expressed coding isoforms contain the catalytic domains of phosphodiesterase 4D (Figure 4). Calcineurin is a widely expressed and highly conserved Ser/Thr phosphatase that binds to phosphodiesterase 4D, inhibiting its proteasomal degradation. Calcineurin in turn is inhibited by tacrolimus (FK506). Therefore, treatment with tacrolimus can be expected to decrease phosphodiesterase 4D activity by promoting its degradation. Our results indicate that treatment with tacrolimus may also shift the distribution of *PDE4D* to isoforms that lack phosphodiesterase activity. *isopret* predicted the same isoform to have the annotation *T cell receptor signaling pathway* (GO:0050852). cAMP is the most potent inhibitor of T-cell activation, and phosphodiesterase 4D is one of the enzymes that degrades cAMP in the cytosol.[29] Therefore, an additional hypothesis that is motivated by our results is that a shift in *PDE4D* isoforms induced by tacrolimus treatment could influence the effects of tacrolimus on T-cell function.

### Isopret-gui

We have implemented the analysis as a Java programming library, command-line tool, and graphical user-interface (GUI) application. The analysis can perform both FET and parent-child overrepresentation analysis with a number of different procedures for multiple testing correction.[2] Splicing patterns and proteins domains are visualized as well as the predictions of GO annotations for for each isoform expressed in the experiment. Isopret-gui is freely available at https://github.com/TheJacksonLaboratory/isopret together with a detailed tutorial. All 100 HBA-DEALS output files analyzed in the work are available for download for use in isopret-gui.

## Discussion

Advances in RNA sequencing technologies have enabled unprecedented accuracy in the quantification of mRNA at the isoform level. However, it remains challenging to assess the functional consequences of differential splicing because, to date, computational resources for GO analysis have largely focused on differential gene expression. GO is a fundamental resource for the biological interpretation of differential expression analysis. Extensive efforts have been made by the bioinformatics community to develop methods that can be used to ascertain the biological “meaning” of experiments by characterizing GO terms that are overrepresented in differentially expressed genes.[2, 30, 31, 32, 33]. However, to our knowledge, no previous method assesses overrepresentation of GO annotations among differentially spliced isoforms. This is presumably due to the paucity of knowledge about isoform-specific functions and the challenges in computational isoform function prediction.

In our view, machine learning approaches for inferring isoform-specific function have three desiderata: (1) they should not rely on supervised learning, which is limited by our current state of knowledge about isoform functions; (2) they should not associate GO terms to genes rather than isoforms; (3) They not make predictions one isoform at a time or one GO term at a time, which precludes information sharing and can result in inconsistencies between different predictions. To our knowledge, no previously published method satisfies these requirements (Supplemental Table S1). In contrast, *isopret* is unsupervised, predicting isoform annotations without using isoform-specific labels for training, learns directly from isoform sequences without using gene elements (e.g. domains), and assigns GO terms to isoforms through a global optimization algorithm, thus avoiding inconsistencies due to local isoform-by-isoform predictions. Our review of 16 published methods found only three that made code or predictions available in a way that enabled us to compare our methods on a comprehensive set of 307 isoform-specific functions derived from our curation of the literature (Supplemental Table S6). Our results show that *isopret* substantially outperformed the other available methods. Additionally, to our knowledge, isopret-gui is the only currently available software for isoform-specific GO overrepresentation analysis.

Our analysis of data from 100 RNA-seq experiments from across the biomedical research spectrum demonstrated that DGE and DAS tend to regulate different sets of GO terms. Thus, a functional analysis of RNA-seq results that focuses solely on differentially expressed genes may miss important functional implications. Our approach to isoform-specific analysis has a number of important limitations that will require future work to address. There remain many technical difficulties in assigning short NGS reads to isoforms[24] that limit the accurate of any downstream algorithms for overrepresentation analysis such as ours. Advances and dropping costs of long-read sequencing technologies may improve accuracy of isoform quantification in the future.[34] It is our hope that methods such as ours will spur more activities in the elucidation of isoform-specific functions, but many community-driven efforts will be required to develop experimental frameworks, databases, and computational methods for isoform-specific analysis.

We provide a Java desktop tool called isopret-gui to analyze RNA-seq experiments for GO overrepresentation for both DGE and DAS. Additional functionality includes an analysis of InterPro domains that are overrepresented in differentially spliced isoforms and visualizations of genes that include visualizations of transcripts, protein domains and the isoform functions predicted by *isopret*. The tool, together with detailed instructions, is freely available at https://github.com/TheJacksonLaboratory/isopret.

## Online Methods

### Data sources

#### Gene Ontology

In order to obtain the Gene Ontology terms associated with each gene, we downloaded the files goa_human.gaf version 2.2 and interpro2GO dated 2020/11/26 from the Gene Ontology website, and the file hgnc_complete_set.txt from the HGNC web site. InterPro domains were obtained from BioMart using the BiomaRt R package. Gene and isoform Ensembl IDs were obtained from Ensembl release 100 human genome version GRCh38 (the file `Homo_sapiens.GRCh38.100.gtf`). GO terms for Molecular Function, Biological Process and Cellular Component were extracted from goa_human.gaf, and combined with the GO terms in interpro2GO. Terms that were common to 10 percent of the genes or more were removed in order to retain terms that correspond to specific functions.

#### Protein-protein interactions

Protein-protein interactions were retrieved from the BioGRID database version 4.4.207 [35]. The PPIs from BioGRID are provided at the gene level, however resources for isoform-isoform interactions that we examined were limited or no longer maintained.

#### Transcript definitions

The Ensembl transcript coordinates were obtained from Ensembl release 100 human genome version GRCh38 from the file `Homo_sapiens.GRCh38.100.gtf`, and the sequences from the corresponding genome file `GRCh38_r91.all.fa`

#### RNA-seq files

We collected RNA-Seq datasets that were deposited to the SRA. These experiments study a broad range of conditions (Supplemental Tables S2-S5), but they all contained two distinct sets of samples that could be compared via a case-control design. The collection contains 109 case-control comparisons as some datasets provide more than one comparison. The mean and median number of case samples was 6 and the median number of control samples was also 6.

### Inferring isoform functions by Expectation Maximization

Our approach is based on the following assumptions: 1) GO terms that are assigned to a gene correspond to a function of one or more isoforms of that gene. 2) The mean pairwise sequence alignment score increases with the number of functions shared by the pair of aligned protein sequences. 3) Given two pairs of isoforms, such that for each pair the genes’ GO terms have at least one common term, and in addition given the GO terms that are assigned to each isoform, the sequence alignment scores of the two pairs are independent random variables.

We briefly introduce some notations that will be used in the description below:

- *G* = *{g*_1_, *g*_2_, …, *g*_*n*_*}* is the set of *n* genes.
- *I* = *{i*_1_, *i*_2_, …, *i*_*p*_*}* is the set of *p* isoforms.
- *GO* = *{t*_1_, *t*_2_, …, *t*_*m*_*}* is the set of *GO* terms, where *t*_*i*_ is an identifier and m the number of terms.
- 2^*GO*^ is the powerset of *GO* terms.
- *𝒯* _*I*_ = *ϕ* : *I →* 2^*GO*^ represents the annotations of the isoforms, e.g. *ϕ*(*i*_1_) are the GO annotations of the isoform *i*_1_
- *𝒯* _*G*_ = *ψ* : *G →* 2^*GO*^ represents the GO annotations of the genes
- 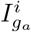 is the isoform *i* belonging to gene *g*_*a*_ (*i ∈ g*_*a*_).
- 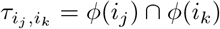 is the set of *GO* terms shared by isoforms *i*_*j*_ and *i*_*k*_.
- *S*(*i*_*j*_, *i*_*k*_) is the sequence similarity between isoforms *i*_*j*_ and *i*_*k*_

The data *S*(*i*_*j*_, *i*_*k*_) can be computed for *i*_*j*_ ∈ *g*_*j*_ and *i*_*k*_ ∈ *g*_*k*_, *j* ≠ *k*, and we suppose (according to our preliminary experimental results) that *S* follows a quadratic relation with respect to the number of GO term annotations shared by *i*_*j*_ and *i*_*k*_:

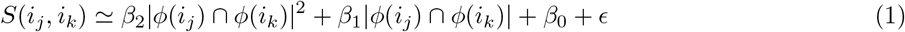

with *β*_0_, *β*_1_ and *β*_2_ unknown parameters. Note that |*ϕ*(*i*_*j*_) *∩ ϕ*(*i*_*k*_)| is the cardinality of GO terms shared by isoforms *i*_*j*_ and *i*_*k*_.

We suppose that *S*(*i*_*j*_, *i*_*k*_) ∼ *N* (*μ*, 1), i.e. that *S* is distributed according to a normal distribution with mean equal to *μ* and *σ* = 1. We suppose that *μ*(*i*_*j*_, *i*_*k*_) = *β*_2_| *ϕ*(*i*_*j*_) ∩ *ϕ*(*i*_*k*_) |^2^ + *β*_1_ |*ϕ*(*i*_*j*_) ∩ *ϕ*(*i*_*k*_) | + *β*_0_. Hence the probability density function of *S* is

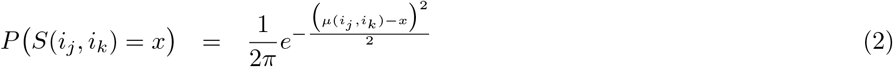

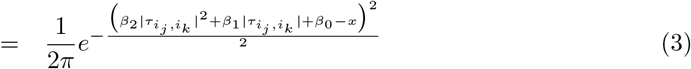

where we use 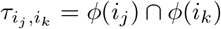 to simplify the notation.

By applying the maximum log-likelihood principle we can maximize the following function:

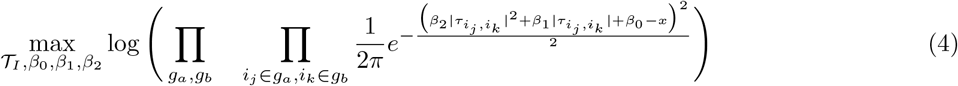

Note that we need to maximize with respect to the isoform annotations *𝒯*_*I*_ and the parameters *β*_0_, *β*_1_, *β*_2_ of the quadratic function, and 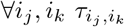 depend on 𝒯_*I*_, i.e. on the annotation of the considered set of isoforms *I*.

Eq. 4 can be better expressed as sums:

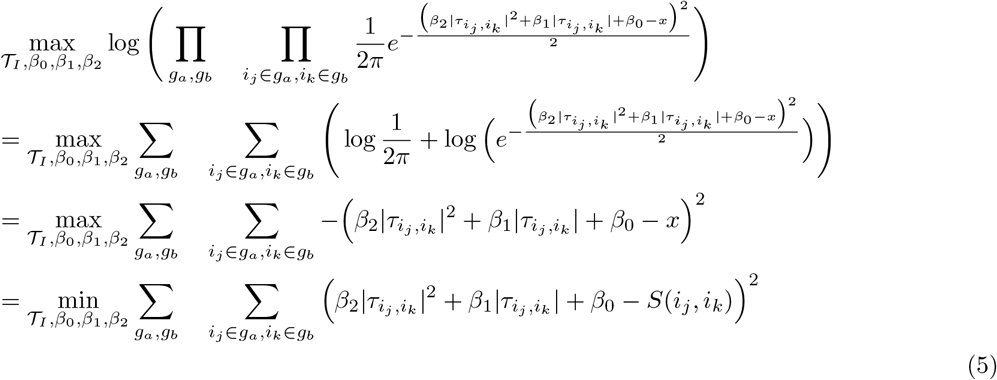

We can optimize eq. 5 through an Expectation Maximization (EM) algorithm:

1. **Initialize** *𝒯*_*I*_ **and model parameters** *β*_0_, *β*_1_, *β*_2_:

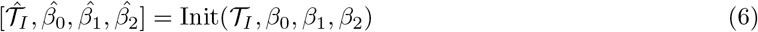
2. **E- step:** Given 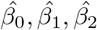:

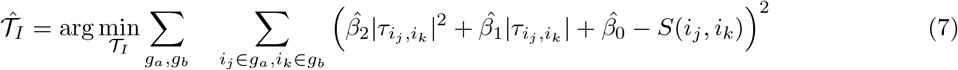
3. **M- step:** Given 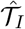 :

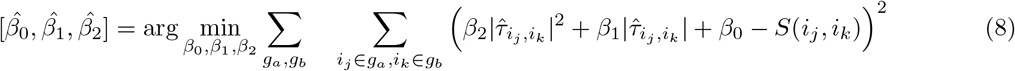
4. **Cycle between E-step ad M-step till to convergence** The final 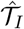 provides the isoform annotations predicted by isopret.

The most computationally demanding part of the EM algorithm is represented by the E-step. Indeed the E-step constitutes an NP-complete problem (Supplemental Note 1). Furthermore, our data contains close to 85,000 isoforms, and consequently computing the sum of log-likelihoods in (5) is a prohibitively expensive operation. This problem is often alleviated in machine learning by the use of mini-batches, namely by repeatedly sampling a subset of the sum and updating the parameters based on that subset [36]. We follow a similar approach by randomly splitting the isoforms into 200 subsets, aligning each pair of isoforms such that they belong to the same subset and their genes share at least one GO term, normalizing the alignment scores and computing an update to the function assignment using a genetic algorithm (GA). In order to estimate the improvement in the total log-likelihood, the log-likelihood sum computed in each subset is divided by the number of pairs of isoforms that are compared in that subset. Convergence of the E step is determined when the change in likelihood over 25 iterations/batches is less than 1/3 (Figure S2). Then, at the M step we compute the least-squares solution to the *β*_*i*_ parameters based on the pairs of isoforms that were compared in the last step of the E-step. For the genetic algorithm (GA) implementation we used the R GA package [37]. The main genetic operator was crossover with a probability of 0.8, with an additional 0.1 probability for mutation. This implementation also applies elitism by default and so the best assignment found is always propagated. The selection probability of a solution was proportional to its rank in increasing order of fitness. As an initial assignment of isoform functions, we used GO terms that were associated with predicted isoform domains, taken from the interpro2GO mapping (Figure S3). The initial guess for *β*_*i*_ parameter values was obtained by fitting the alignment scores from the subset of isoforms in the first split to the number of common GO terms in the initial assignment. Since the three subsets of GO, Molecular Function, Biological Process and Cellular Component have synonymous terms, we performed the optimization separately for the union of the interpro2GO mapping and each one of the three GO subsets. We reasoned that Cellular Component terms would not be able to account for the same degree of sequence diversity as Biological Process and Molecular Function, and therefore when running the optimization for this subset we started from the assignment of GO terms that were assigned in the optimization of Biological Process. The sequence alignment is computed using the Smith-Waterman local alignment algorithm implemented in the pairwiseAlignment function in R’s Biostrings package, with the BLOSUM62 substitution matrix and all other parameters taking their default values. In this work, we included only protein-coding isoforms, although a generalization that aligns nucleotide sequence instead would be conceptually similar. Normalization of alignment scores was performed using the *log_x* function of the bestNormalize R package [38] with default parameters.

### Evaluation

We reviewed the literature for papers that determine the function of isoform that can be mapped to Ensembl IDs and whose function can be mapped to GO terms. We read original publications cited in a systematic evaluation of the literature on alternative splicing[8], and splicing events were included if a GO annotation could be chosen to represent the documentation differential function of an isoform. In some cases, differential functions reported in this work represented phenomena such as increased proliferation that were related to cancer and could not be assigned GO terms unambiguously. These were not included in our curation. The result was a collection of 307 examples where an isoform was shown to be either associated with a certain GO term or not associated with it (Supplemental Table S6). As a comparison for Isopret’s performance, we used three other GO term assignment methods: (1) For each isoform, assign the GO terms that interpro2GO maps to its InterPro domains (2) The IsoResolve [17] tool. IsoResolve uses gene expression profiles to make predictions, and the tool comes with a relatively small expression dataset for mouse. In order to use it for human, we used the Lung dataset transcript-per-million counts from the Genotype-Tissue Expression (GTEx) project.[39] (3) The DIFFUSE tool [16] that applies a deep learning approach to isoform function prediction. We used the prediction of GO terms for human that are provided with the tool. We also tried to use the predictions provided by FINER [40], however none of them matched entries in our collection.

### Mapping of RNA-seq data and isoform calling

For the the analysis of the collection of 100 RNA Sequencing datasets, for each case vs control comparison, RNA-seq data were downloaded from the NCBI Sequence Read Archive (SRA) resource [41]. We have selected a diverse set of conditions in order to obtain a diverse set of enriched GO terms (Figure S5). All datasets were downloaded and processed using a snakemake pipeline that performs the following steps: samples are downloaded from the SRA, quality-control using fastp [42], alignment to Genome Reference Consortium Human Build 38 version 91 using STAR [43], and isoform quantification by RSEM [44] and TMM normalization [45]. For calling differentially-expressed genes and differentially-spliced isoforms we use HBA-DEALS [19, 46].

### Isoform-focused Gene Ontology overrepresentation analysis

We implemented GO overrepresentation analysis with a modernized (Java 17) version of the Ontologizer software.[47, 2] The Fischer-exact test (term for term), and parent-child union and intersection methods and several multiple testing correction methods were implemented.[18] For gene-level (standard) overrepresentation analysis, the population set is taken to be the set of all genes with at least one read in the RNA-seq experiment, and the study set is taken to be the set of differentially expressed genes. The standard, gene-level GO annotations are used for this analysis.

For isoform-level analysis, the population set is taken to be the set of all isoforms with at least one read in the RNA-seq experiment, and the study set is taken to be the set of differentially spliced isoforms. The GO annotations inferred by *isopret* are used for this analysis.

### InterPro domain overrepresentation analysis

Using the Fischer-exact test overrepresentation software mentioned in the previous section, an overrepresentation test is performed by defining isoforms with the presence of one or more ProSite domain annotations to be annotated to the ProSite entry. The population set is taken to be the set of all isoforms with at least one read in the RNA-seq experiment, and the study set is taken to be the set of differentially spliced isoforms. The GO annotations inferred by *isopret* are used for this analysis. Multiple-testing correction was performed with the Bonferroni procedure.

## Acknowledgements

The authors thank Jane Cha for designing Figure 1.

## Author contributions

G.K. and P.N.R. conceived the project. G.K. performed all computational analyses. G.K. and P.N.R. developed the computer code to implement the algorithms presented in the manuscript. L.C., J.C.S., G.K., and P.N.R. performed biocuration of functional annotations of isoforms. E.C., P.H., J.R., C.J.M., and G.V. contributed to the presentation of the algorithm and the interpretation of the results. P.N.R. supervised the project. G.K. and P.N.R. wrote the manuscript with contributions from all authors.

## Supplementary information

**Supplementary Information** Supplementary Figs. S1-S6, Tables S1-S5, and Supplementary Note 1

**Supplementary Table** Supplementary Table S6 with curated isoform function predictions

