## Supplemental Figures, Tables, and Note for "An algorithmic framework for isoform-specific functional analysis"

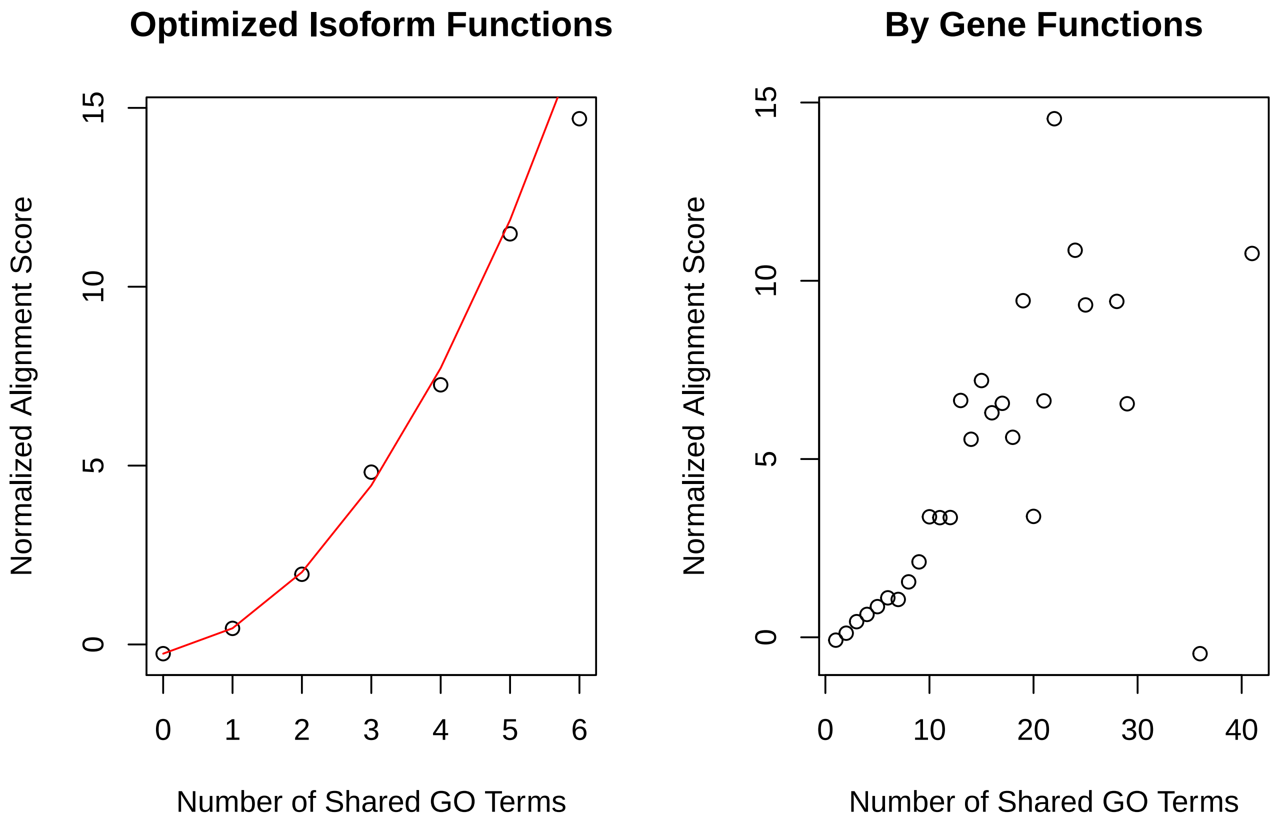

**Figure S1: Mean sequence alignment score as a function of the number of shared domains.** In order to alleviate the computational cost of the E step of the EM algorithm, we repeatedly split the isoforms into 200 random subsets and run a genetic algorithm in each subset for a fixed number of iterations. Here, for the last partition into 200 isoform sets, mean normalized sequence alignment scores were plotted against the number of GO terms shared by pairs of isoforms (left) and the number of GO terms shared by the pairs of genes that contain the isoforms (right). The red line in the left frame corresponds to the quadratic model (expressed in equation (5) in the main text) using the final $\beta_i$ values. The figure was generated for the optimization of GO Molecular Function+Interpro2GO.

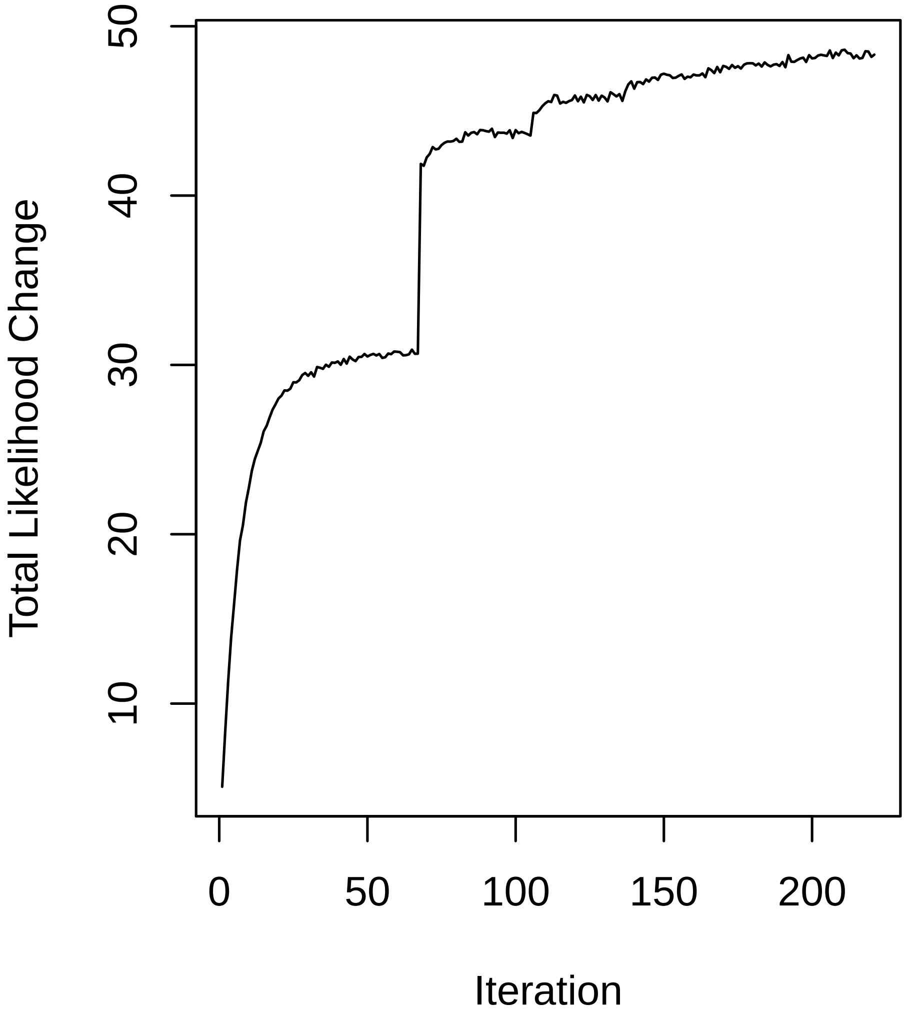

**Figure S2: Changes in likelihood over the course of the algorithm.** Executing the E step of the algorithm is computationally challenging, and therefore we repeatedly split the isoforms into 200 random subsets and use them to guide the search instead of the full log-likelihood. Here, a value on the x-axis corresponds to one optimization step, i.e. a random partition of all isoforms into 200 sets and optimization of the GO term assignments within each set. The y-axis shows the sum of likelihood changes divided by the number of log-likelihood terms over all 200 sets, starting from the difference between the value of the objective after the second step and its value after the first step. The E step terminates when the sum of changes over its last 25 partitions does not exceed a small threshold, after which the M step optimizes the parameters that map the number of shared GO terms to the normalized alignment score. The figure was generated for the optimization of GO Molecular Function+Interpro2GO.

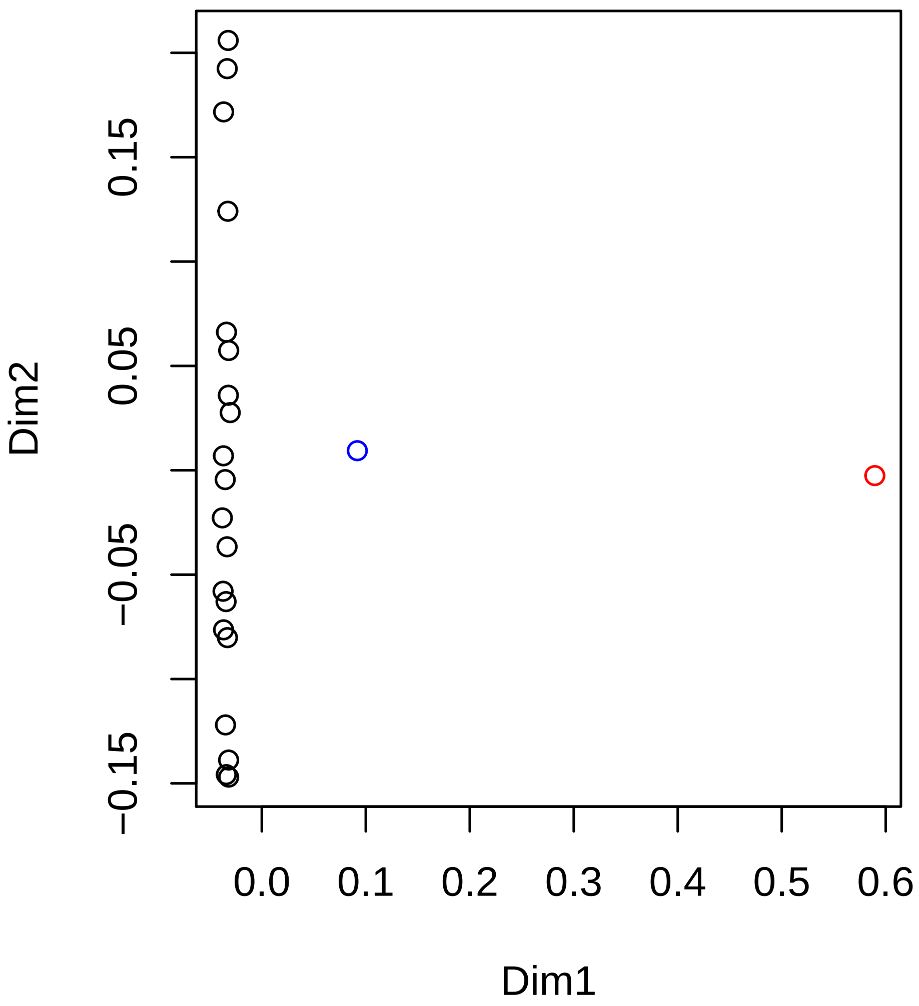

**Figure S3: Multidimensional scaling of the initial vector of isoform function assignment, the final vector after convergence, and random permutations of the latter vector.** Each assignment of GO functions to all isoforms corresponds to a binary vector, with a value of 1 in entries that correspond to a GO term that was selected for an isoform and 0 for a GO term that was not. Different assignments/Boolean vectors can be visualized using multidimensional scaling. Here, the blue dot correspond to the optimal assignment, the red dot to the initial assignment that is based on Interpro2GO, and the black dots to random permutations of the optimal assignment. The distance measure used is the binary distance implemented in R's dist function. As illustrated in the figure, randomly permuting the optimal assignment does result in a permutation that is closer to the initial assignment, which implies that they choose disjoint sets of function more than expected by chance, or that one of them consistently chooses more functions, or both. The figure was generated for the optimization of GO Molecular Function+Interpro2GO.

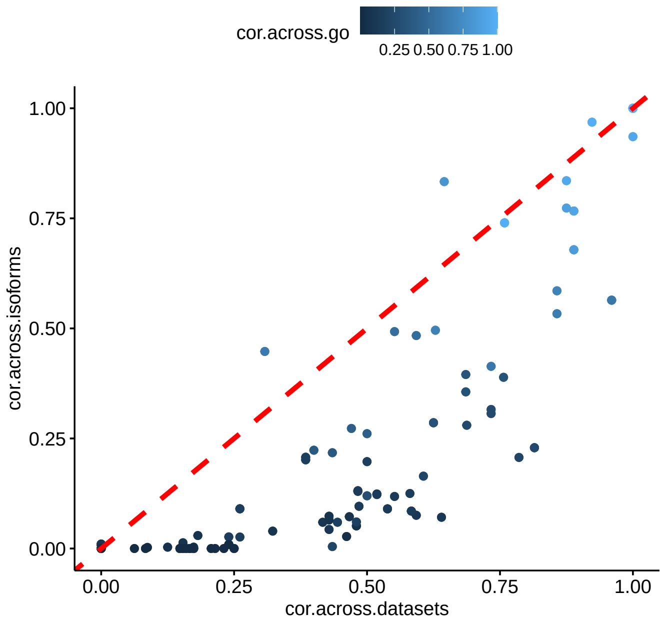

**Figure S4: Co-occurrence of Interpro domains in datasets and on isoforms.** The value on the x-axis of the scatter plot corresponds to the number of common datasets that a pair of Interpro domains is enriched in, divided by the total number of datasets that either of them is enriched in. The value on the y-axis corresponds to the number of transcripts that a pair of domains appears on together divided by the number of transcripts either of them appears on. Each dot on the scatter plot corresponds to one pair of Interpro domains. Only domains that are DAS-enriched in at least 10 datasets are plotted. The dashed red line is y=x. The figure shows that for domains that occur in at least 10 datasets, co-enrichment is correlated with co-occurrence (Pearson correlation 0.85), where the former is higher than the latter, likely since co-occurrence is associated with a functional relationship that can be carried out by two different isoforms, each carrying only one of the individual domains.

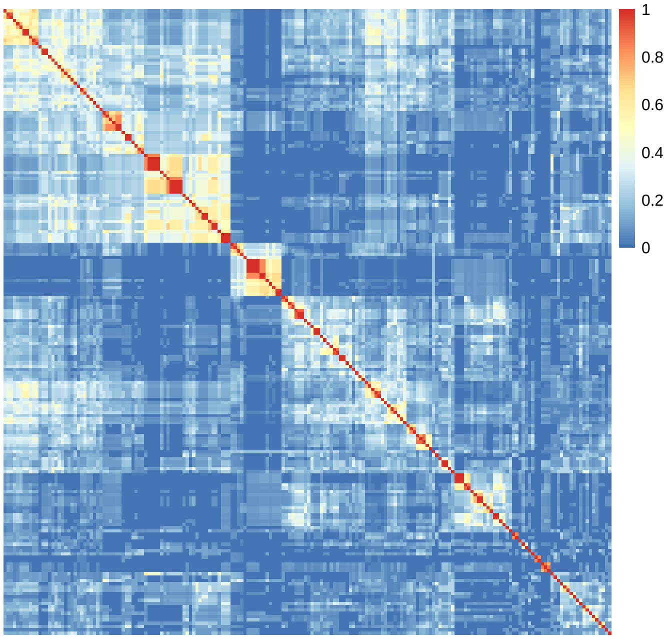

**Figure S5: Co-occurrence of GO terms across datasets** The heatmap shows clusters of GO terms that co-occur as DAS-enriched in the same datasets, using our collection of 100 RNA-Seq datasets. The rows and columns correspond to GO terms that are enriched in at least 5 datasets, and the color corresponds to the Jaccard index between datasets in which they are DAS-enriched, i.e. the proportion of datasets in which both GO terms are DAS-enriched, out of all datasets that have either GO term as DAS-enriched. The figure shows that there are some clusters of datasets that are related enrichment-wise, although the overlap in DAS enrichment over all the collection is relatively low, indicating a diverse set of differential-splicing responses.

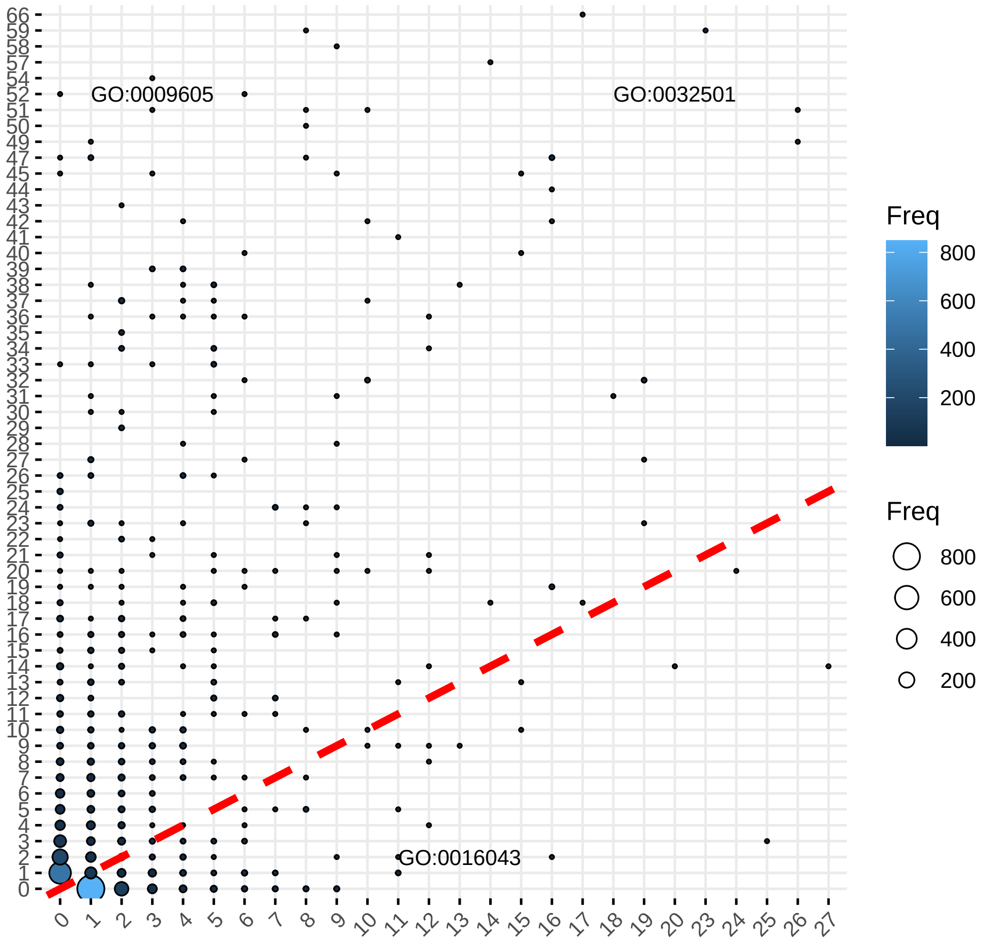

**Figure S6: DGE vs DAS counts for all GOP terms.** The plot illustrates the number of datasets in which every GO term is DAS- and DGE-enriched, for GO terms that are enriched in at least one dataset in our collection. The x-coordinate corresponds to the number of datasets in which a GO term is enriched for DAS, the y-coordinate corresponds to the number of datasets in which a GO term is enriched for DGE, and the size of the dot corresponds to the number of GO terms for the corresponding x and y axis values apply. Some of the GO terms are displayed next to the dots that represent them. The term GO:0032501 (multicellular organismal process is enriched ) appears in many datasets as DGE and in many as DAS, whereas the terms GO:0016043 (cellular component organization) and GO:0009605 (response to external stimulus) are mostly enriched for only one mechanism, DAS or DGE, respectively.

| Author | Year | Ref. | Algorithm name | Predictions? | Executable? | Tested? |
|---|---|---|---|---|---|---|
| Eski | 2013 | [1] | IsoPred | - | - | - |
| Li | 2014 | [2] | IsoFP | - | - | - |
| Li | 2014 | [3] | not named | - | - | - |
| Tseng | 2015 | [4] | IIIDB | - | - | - |
| Mitchell | 2015 | [5] | interpro2go | ✓ | - | ✓ |
| Luo | 2017 | [6] | - | - | - | |
| Shaw | 2018 | [7] | DeepIsoFun | - | - | * |
| Chen | 2019 | [8] | DIFFUSE | ✓ | - | ✓ |
| Ferrer-Bonsoms | 2020 | [9] | IsoGO | - | - | - |
| Yu | 2019 | [10] | IsoFun | - | ** | - |
| Yunes | 2018 | [11] | Effusion | - | *** | - |
| Li | 2020 | [12] | IsoResolve | - | ✓ | ✓ |
| Wang | 2020 | [13] | Diso-Fun | - | - | - |
| Yu | 2021 | [14] | TS-Isofun | - | - | - |
| Yu | 2021 | [15] | DMIL-IsoFun | - | - | - |
| Chen | 2021 | [16] | FINER | ✓ | - | ✓ |

**Table S1: Availability of predictions or executable code of previous approaches to isoform prediction**. The table shows published algorithms and indicated whether predictions made by the algorithm, are available ("Predictions?") or whether script or program is available which could be used to generate predictions for the algorithm ("Executable?"). We followed links from the original publications and searched for updated links using standard internet search engines. In some cases, following the original links produces a 404 Page Not Found error (e.g., [1, 2]. In others, the original papers did not provide predictions or code (e.g.,[6]). (*) We did not test DeepIsoFun, because is was presented by the same group as DIFFUSE, which is a later paper and was reported to outperform DeepIsoFun. (**) IsoFun, Diso-Fun and DMIL-IsoFun require a license to matlab[10, 15, 13]; FINER provided predictions for 471 GO terms but none of its predictions matched entries in our gold standard [16]. No open-source version is available. (***) Unable to run provided code.

| cohort | case:control | category | author | year | pmid |
|---|---|---|---|---|---|
| SRP120040_105 | 5:4 | neurodegenerative disease | Osipovitch M | 2019 | 30554964 |
| SRP120040_106 | 7:4 | neurodegenerative disease | Osipovitch M | 2019 | 30554964 |
| SRP188206_107 | 12:12 | Neurodegenerative disease | Lavin KM | 2020 | 31829804 |
| SRP188206_108 | 12:12 | Neurodegenerative disease | Lavin KM | 2020 | 31829804 |
| SRP051518_111 | 4:7 | Kawasaki disease | Rowley AH | 2015 | 26679344 |
| SRP051518_112 | 4:7 | Kawasaki disease | Rowley AH | 2015 | 26679344 |
| SRP150872_114 | 28:32 | lupus | Davenport EE | 2018 | 30340504 |
| SRP332552_117 | 12:12 | diabetes | Gabriel BM | 2021 | 34669477 |
| SRP223448_119 | 10:22 | schizophrenia | Perez JM | 2021 | 32152472 |
| SRP325897_221 | 10:12 | liver pathophysiology | Lv Y | 2021 | 34413880 |
| SRP228563_222 | 11:12 | sickle cell disease | Creary S | 2020 | 32487996 |
| SRP318393_244 | 10:10 | Sjögren | Verstappen GM | 2021 | 34295332 |
| SRP287301_247 | 8:13 | enteric dysfunction. | Haberman Y | 2021 | 33524399 |
| SRP144583_251 | 18:38 | enteric fever | Blohmke CJ | 2019 | 31468702 |
| SRP233184_261 | 12:12 | aortic aneurysm | Chen PY | 2020 | 32243809 |
| SRP222046_263 | 4:4 | abdominal aortic aneurysm | Lu H | 2020 | 32354235 |
| SRP219588_264 | 18:9 | Vibrio vulnificus | Kim BS | 2020 | 32817457 |
| SRP279451_265 | 8:8 | atopic dermatitis | Möbus L | 2021 | 32615169 |
| SRP165679_266 | 6:38 | psoriasis | Tsoi LC | 2019 | 30641038 |
| SRP289501_267 | 9:9 | asthma | Helling BA | 2020 | 33188283 |
| SRP234559_281 | 12:8 | idiopathic aplastic anemia | Lim SP | 2020 | 32294156 |
| SRP335222_286 | 18:12 | basophil physiology | Puan KJ | 2021 | 34215830 |
| SRP292025_287 | 6:11 | neurodegenerative disease | Garofalo M | 2020 | 33327559 |
| SRP292025_288 | 6:11 | neurodegenerative disease | Garofalo M | 2020 | 33327559 |
| SRP346219_294 | 4:4 | peripheral vascular disease | Gross DA | 2022 | 34793334 |
| SRP258588_299 | 12:24 | Burn-McKeown syndrome | Wood KA | 2020 | 32735620 |
| SRP262152_304 | 4:4 | atherosclerosis | Kim JB | 2020 | 32441123 |
| SRP305908_306 | 4:4 | intracerebral hemorrhage | Goods BA | 2021 | 33749664 |
| SRP040622_308 | 7:6 | stroke | Huttner HB | 2014 | 24747576 |
| SRP328165_310 | 5:8 | liver pathophysiology | Zhang IW | 2022 | 34450236 |
| SRP191103_312 | 8:12 | gastroparesis | Herring BP | 2019 | 31221130 |
| SRP279451_316 | 21:21 | atopic dermatitis | Möbus L | 2021 | 32615169 |
| SRP161727_321 | 9:5 | Diamond-Blackfan anemia | Ulirsch JC | 2018 | 30503522 |
| SRP134188_322 | 8:10 | dystrophic epidermolysis bullosa | Cho RJ | 2018 | 30135250 |
| SRP272540_323 | 8:6 | immune thrombocytopenia | Han P | 2021 | 33876188 |
| SRP189352_327 | 4:15 | SF3B1 mutations | Zhang J | 2019 | 31474574 |
| SRP342557_328 | 5:5 | epilepsy | Gomes-Duarte A | 2022 | 35310884 |

**Table S2:** 33 disease-related cohorts analyzed in this work. Colummns: **SRP**: Project (study) accession number in NCBI's Sequence Read Archive [17]; **case:control**: RNA-seq sample counts for cases and controls; **category**: representative topic addressed by the RNA-seq experiment; **author** First author of the publication associated with the project; **year**: year of publication; **pmid**: PubMed identifier.

| cohort | case:control | category | author | year | pmid |
|---|---|---|---|---|---|
| SRP162188_233 | 9:18 | DRUG-seq | Ye C | 2018 | 30333485 |
| SRP219481_236 | 6:6 | erythropoiesis | Rossmann MP | 2021 | 33986176 |
| SRP117167_242 | 7:7 | memory T cell physiology | Belarif L | 2018 | 30367166 |
| SRP258620_243 | 7:5 | macrophage physiology | Gutbier S | 2020 | 32645954 |
| SRP218229_249 | 12:8 | adipose tissue | Vijay J | 2020 | 32066997 |
| SRP218230_253 | 10:6 | adipose tissue | Vijay J | 2020 | 32066997 |
| SRP260413_255 | 4:6 | radiation damage | Brambilla F | 2020 | 32710624 |
| SRP302877_269 | 4:4 | proteasome | Cáceres-Gutiérrez RE | 2022 | 34739170 |
| SRP229996_271 | 8:4 | CD8 T cell physiology | Jansen CS | 2019 | 31827286 |
| SRP100829_272 | 5:5 | oocyte physiology | Reyes JM | 2017 | 29025019 |
| SRP202034_274 | 5:4 | liver physiology | Mendoza A | 2021 | 34784250 |
| SRP224022_275 | 6:6 | intestinal enteroids | Chang-Graham AL | 2019 | 31029854 |
| SRP221491_282 | 4:4 | basal cell-like cells | Lu J | 2021 | 32365352 |
| SRP229589_283 | 4:5 | tacrolimus | Dai C | 2020 | 31941840 |
| SRP168076_285 | 6:6 | basophil physiology | Puan KJ | 2021 | 34215830 |
| SRP335222_286 | 18:12 | basophil physiology | Puan KJ | 2021 | 34215830 |
| SRP094851_293 | 4:4 | cardiomyocyte physiology | Necela BM | 2017 | 28101782 |
| SRP300738_295 | 5:5 | oocyte physiology | Ntostis P | 2021 | 34755188 |
| SRP257383_296 | 4:4 | development | Valcourt JR | 2021 | 34758327 |
| SRP344260_297 | 4:4 | pericyte physiology | Rezaei-Lotfi S | 2021 | 34886891 |
| SRP253111_303 | 6:6 | colon physiology | Bergenheim F | 2020 | 32891909 |
| SRP162681_307 | 4:4 | cell cycle | Mahmoud AD | 2019 | 31339449 |
| SRP297875_311 | 6:6 | gene expression | Grundman J | 2021 | 34587152 |
| SRP255876_313 | 8:8 | exposure to formaldehyde | Gonzalez-Rivera JC | 2020 | 33024153 |
| SRP334204_317 | 4:4 | macrophage physiology | De M | 2022 | 35115664 |
| SRP334204_318 | 4:4 | macrophage physiology | De M | 2022 | 35115664 |
| SRP334204_319 | 4:4 | macrophage physiology | De M | 2022 | 35115664 |
| SRP334204_320 | 4:4 | macrophage physiology | De M | 2022 | 35115664 |
| SRP217536_324 | 9:10 | high-protein diet | Xu C | 2020 | 32652799 |
| SRP149366_329 | 4:4 | breast cells | Meng P | 2019 | 30993572 |

**Table S3:** 28 RNA-seq experiments devoted primarily to the study of physiology, gene expression or cell biology. Columms: see Table **S2**.

| cohort | case:control | category | author | year | pmid |
|---|---|---|---|---|---|
| SRP219837_113 | 7:5 | colorectal cancer | Orouji E | 2022 | 3405950 |
| SRP065445_115 | 7:5 | Histiocytic neoplasms | Diamond EL | 2016 | 2656687 |
| SRP286904_223 | 8:8 | AML | Ho JM | 2020 | 3314733 |
| SRP090124_224 | 26:11 | breast cancer | Pouliot MC | 2017 | 2910825 |
| SRP215936_225 | 13:18 | breast cancer | Arruabarrena-Aristorena A | 2020 | 3288843 |
| SRP042620_226 | 42:30 | breast cancer | Varley KE | 2014 | 2492967 |
| SRP281892_227 | 52:26 | melanoma | Hong X | 2021 | 3320373 |
| SRP090849_228 | 22:9 | osteosarcoma | Scott MC | 2018 | 2906651 |
| SRP026537_229 | 7:15 | breast cancer | Daemen A | 2013 | 2417611 |
| SRP233497_230 | 21:6 | pancreatic cancer | Porter RL | 2019 | 3184392 |
| SRP331153_231 | 4:4 | breast cancer | Arruabarrena-Aristorena A | 2020 | 3288843 |
| SRP092413_235 | 26:10 | neuroblastoma | Harenza JL | 2017 | 2835038 |
| SRP217026_237 | 6:6 | pancreatic cancer | Salvador-Barbero B | 2020 | 3210937 |
| SRP134389_238 | 18:4 | breast cancer | Ye IC | 2018 | 3003785 |
| SRP247679_239 | 8:12 | cancer | Pearson JD | 2021 | 3427092 |
| SRP119676_240 | 32:30 | liver cancer | Hooks KB | 2018 | 2915277 |
| SRP111914_241 | 10:19 | liver cancer | Li S | 2019 | 3001461 |
| SRP050440_246 | 7:6 | resistance to BET inhibition | Rathert P | 2015 | 2636779 |
| SRP220467_248 | 8:6 | retinoblastoma | Liu H | 2020 | 3331819 |
| SRP313282_254 | 9:11 | lung cancer | Zhang T | 2021 | 3449386 |
| SRP012167_256 | 5:4 | parathyroid adenoma | Haglund F | 2012 | 2302418 |
| SRP332697_257 | 6:6 | parathyroid adenoma | Haglund F | 2012 | 2302418 |
| SRP332697_257 | 6:6 | parathyroid adenoma | Haglund F | 2012 | 2302418 |
| SRP301216_258 | 5:5 | colorectal cancer | Hong Q | 2021 | 3445814 |
| SRP327911_260 | 5:5 | malignant pleomorphic adenoma | Han Z | 2022 | 3498685 |
| SRP278517_270 | 11:6 | ovarian cancer | Cardillo N | 2021 | 3349912 |
| SRP303687_273 | 7:19 | thyroid cancer | He H | 2021 | 3423898 |
| SRP302218_277 | 4:4 | sarcoma | Carrabotta M | 2022 | 3490360 |
| SRP276412_278 | 5:5 | GI stromal | Shao Y | 2021 | 3445801 |
| SRP183071_280 | 7:9 | DLBCL | McCord R | 2019 | 3077036 |
| SRP311634_284 | 10:10 | pancreatic cancer | Farshadi EA | 2021 | 3458011 |
| SRP183757_290 | 5:5 | EMT-chemoresistance | Sale MJ | 2019 | 3104868 |
| SRP267712_291 | 6:6 | melanoma | Grigore F | 2020 | 3262917 |
| SRP336449_292 | 5:12 | head and neck cancer | Bouhaddou M | 2021 | 3454697 |
| SRP253895_301 | 6:6 | cancer | Chan TW | 2020 | 3310617 |
| SRP253895_302 | 6:6 | cancer | Chan TW | 2020 | 3310617 |
| SRP226592_305 | 6:6 | resistance phenotypes | Johnson AG | 2020 | 3246344 |
| SRP312693_314 | 4:4 | medulloblastoma | Rea J | 2021 | 3435975 |
| SRP254646_315 | 11:11 | prostate cancer | He YD | 2021 | 3430126 |

**Table S4:** 39 RNA-seq experiments devoted primarily to the study of cancer. Columms: see Table **S2**.

| cohort | case:control | category | author | year | pmid |
|---|---|---|---|---|---|
| SRP049605_116 | 28:13 | Lyme disease | Bouquet J | 2016 | 26873097 |
| SRP286302_118 | 6:6 | Trypanosoma cruzi | Gil-Jaramillo N | 2021 | 33897690 |
| SRP095674_220 | 4:6 | Schistosoma haematobium | Labuda LA | 2020 | 31844885 |
| SRP134018_250 | 15:24 | Tuberculous meningitis | Rohlwink UK | 2019 | 31434901 |
| SRP320156_309 | 6:6 | tuberculosis | Reichmann MT | 2021 | 34128839 |

**Table S5:** 5 RNA-seq experiments related to infectious disease. Columms: see Table **S2**.

### Supplementary Note 1

A reduction is a method used in theoretical computer science to transform one problem into another problem. In this section, we show that the graph 3-coloring problem, which is NP-complete, can be transformed into the E step of the isoform function assignment problem as posed in the main manuscript (we will call it isoform-GO-assignment for brevity). Although it remains to be proved, it is widely believed that no polynomial time algorithms exist for finding solutions to NP-complete problems. Colloquially speaking NP-complete problems belong to a class of problems that are difficult to solve efficiently. The purpose of this proof is to motivate the need for a heuristic (approximation) at the E-step of the EM algorithm described in the main text.

Given a graph $G(V, E)$ where $V$ is the set of vertices and $E \subset V \times V$ is the set of edges, a $k$-coloring assigns to each node $v \in V$ a label $l_v \in 1, 2, \ldots, k$ such that if $(u, v) \in E$ then $l_v \neq l_u$.

Let $G(V, E)$ be an instance of a 3-coloring, i.e. any input to the 3-coloring problem. We perform the following polynomial-time construction of an instance of isoform-GO-assignment:

1. For each node $v \in V$ we create one isoform.
2. Assign the isoforms to genes arbitrarily. Every gene has the same set of 3 GO terms.
3. The observed sequence similarity scores will be $S(i, j) = \begin{cases} -1 & (i, j) \in E \\ 0 & (i, j) \notin E \end{cases}$
4. Let $\beta_0 = -1, \beta_1 = 2, \beta_2 = 0$, and define a quadratic equation to predict the observed sequence score as a function of the number of shared GO terms, $f(n) = \beta_0 + \beta_1 n + \beta_2 n^2$.

**Claim 1.** *There is a 3-coloring for the graph if and only if there is an isoform-GO-assignment where the sum of absolute differences between predicted and observed sequence similarities is* $\binom{|V|}{2} - |E|$ *, i.e. the sum is equal to the number of vertex pairs that do not have an edge between them.*

*Proof.* First direction: Assume that there is a 3-coloring for $G$. We assign GO terms as follows: For each isoform $i$ assign the GO term with the index of the color of its corresponding node in the graph. Since nodes that have an edge between them do not share a color, the corresponding isoforms will not share a GO term, and the predicted sequence similarity between them will be $\beta_0 = -1$. By the construction this is exactly the observed sequence similarity score, so the sum of differences for these nodes will be 0. For nodes that do not have an edge between, by the construction they can either share one GO term or zero GO terms. So the predicted sequence similarity score will be either 1 or -1, in both cases an absolute difference of 1 from the observed sequence similarity of 0. The total sum of absolute differences is then $\binom{|V|}{2} - |E|$, which is the number of these nodes.

Second direction: Now assume that there is an isoform function assignment such that the sum of absolute differences between predicted and observed sequence similarities is $\binom{|V|}{2} - |E|$. First, we will show that the sum of absolute differences between isoforms $i$ and $j$ such that $(i, j) \notin E$ is at least $\binom{|V|}{2} - |E|$:

If $i$ and $j$ share zero or one GO terms, then as we have seen in the other direction of the proof, the absolute difference between the predicted and observed sequence similarity is 1. If they share two GO terms, then the predicted similarity is $\beta_0 + \beta_1 \cdot 2 + \beta_2 \cdot 2^2 = 3$, and the absolute difference is $|3 - 0| = 3$. Similarly, if they share three GO terms the absolute difference is 5. Since as we have shown the minimal absolute difference is 1, and since there are $\binom{|V|}{2} - |E|$ such isoform pairs, the sum of absolute differences between predicted and observed sequence similarity scores for these isoform pairs is at least $\binom{|V|}{2} - |E|$.

Since absolute difference are non-negative, and since the total sum of absolute difference in the solution is $\binom{|V|}{2} - |E|$, all other differences must be 0. Since the sequence similarity between all other pairs of isoforms, i.e. those for which $(i,j) \in E$ is -1, this is also the predicted sequence similarity scores between them, and this can only happen if they do not share a GO term. Now, assign to each node the color that corresponds to the index of the GO term that was assigned to its corresponding isoform. If the isoform was assigned more than one G0 term, arbitrarily select one term/color from those that were assigned to it. By the construction, none of the adjacent nodes will be assigned the same color. This completes the proof. □

**Notes**

1. The reduction can be slightly changed such that isoforms can be left without any GO term assigned to them in the definition of the GO assignment problem. To obtain this, we connect each node/isoform to $|E|$ new nodes that are connected only to it, and have sequence similarity 1 to it - then it is easy to see that in the optimal solution each isoform is assigned a GO term.

2. We used the $L_1$ norm for difference between predicted and observed sequence similarities in the definition of the GO assignment problem, but the same reduction can be done with the $L_2$ norm with minimal changes.

[5] Alex Mitchell, Hsin-Yu Chang, Louise Daugherty, Matthew Fraser, Sarah Hunter, Rodrigo Lopez, Craig McAnulla, Conor McMenamin, Gift Nuka, Sebastien Pesseat, Amaia Sangrador-Vegas, Maxim Scheremetjew, Claudia Rato, Siew-Yit Yong, Alex Bateman, Marco Punta, Teresa K. Attwood, Christian J. A. Sigrist, Nicole Redaschi, Catherine Rivoire, Ioannis Xenarios, Daniel Kahn, Dominique Guyot, Peer Bork, Ivica Letunic, Julian Gough, Matt Oates, Daniel Haft, Hongzhan Huang, Darren A. Natale, Cathy H. Wu, Christine Orengo, Ian Sillitoe, Huaiyu Mi, Paul D. Thomas, and Robert D. Finn. The InterPro protein families database: the classification resource after 15 years. *Nucleic acids research*, 43:D213–D221, January 2015.

[13] Keyao Wang, Jun Wang, Carlotta Domeniconi, Xiangliang Zhang, and Guoxian Yu. Differentiating isoform functions with collaborative matrix factorization. *Bioinformatics*, November 2019.

[14] Guoxian Yu, Qiuyue Huang, Xiangliang Zhang, Maozu Guo, and Jun Wang. Tissue specificity based isoform function prediction. *IEEE/ACM transactions on computational biology and bioinformatics*, PP, June 2021.

[15] Guoxian Yu, Guangjie Zhou, Xiangliang Zhang, Carlotta Domeniconi, and Maozu Guo. DMIL-IsoFun: predicting isoform function using deep multi-instance learning. *Bioinformatics (Oxford, England)*, July 2021.

[16] Hao Chen, Dipan Shaw, Dongbo Bu, and Tao Jiang. FINER: enhancing the prediction of tissue-specific functions of isoforms by refining isoform interaction networks. *NAR genomics and bioinformatics*, 3:lqab057, June 2021.

[17] Rasko Leinonen, Hideaki Sugawara, Martin Shumway, and International Nucleotide Sequence Database Collaboration. The sequence read archive. *Nucleic acids research*, 39:D19–D21, January 2011.
